# Tetravalent SARS-CoV-2 S1 Subunit Protein Vaccination Elicits Robust Humoral and Cellular Immune Responses in SIV-Infected Rhesus Macaque Controllers

**DOI:** 10.1101/2023.03.15.532808

**Authors:** Muhammad S. Khan, Eun Kim, Quentin Le Hingrat, Adam Kleinman, Alessandro Ferrari, Jose C Sammartino, Elena Percivalle, Cuiling Xu, Shaohua Huang, Thomas W. Kenniston, Irene Cassaniti, Fausto Baldanti, Ivona Pandrea, Andrea Gambotto, Cristian Apetrei

**Author notes:** **Corresponding Authors:** Andrea Gambotto, MD, Department of Surgery, University of Pittsburgh School of Medicine, W1148 Biomedical Science Tower, 200 Lothrop St. Pittsburgh, PA 15261, USA, Cristian Apetrei, MD, PhD, Division of Infectious Diseases, Department of Medicine, University of Pittsburgh School of Medicine, 3550 Terrace St., Pittsburgh, PA 15261, USA. Authors Contributed Equally (Muhammad S. Khan, Eun Kim).

## Abstract

The COVID-19 pandemic has highlighted the need for safe and effective vaccines to be rapidly developed and distributed worldwide, especially considering the emergence of new SARS-CoV-2 variants. Protein subunit vaccines have emerged as a promising approach due to their proven safety record and ability to elicit robust immune responses. In this study, we evaluated the immunogenicity and efficacy of an adjuvanted tetravalent S1 subunit protein COVID-19 vaccine candidate composed of the Wuhan, B.1.1.7 variant, B.1.351 variant, and P.1 variant spike proteins in a nonhuman primate model with controlled SIVsab infection. The vaccine candidate induced both humoral and cellular immune responses, with T- and B cell responses mainly peaking post-boost immunization. The vaccine also elicited neutralizing and cross-reactive antibodies, ACE2 blocking antibodies, and T-cell responses, including spike specific CD4^+^ T cells. Importantly, the vaccine candidate was able to generate Omicron variant spike binding and ACE2 blocking antibodies without specifically vaccinating with Omicron, suggesting potential broad protection against emerging variants. The tetravalent composition of the vaccine candidate has significant implications for COVID-19 vaccine development and implementation, providing broad antibody responses against numerous SARS-CoV-2 variants.

## Introduction

The coronavirus disease 2019 (COVID-19) pandemic caused by the severe acute respiratory syndrome coronavirus 2 (SARS-CoV-2) has had an unprecedented impact on global health, economy, and society. The COVID-19 pandemic consisted of over 675 million cases, with 6.5 million deaths, and 13 billion COVID-19 vaccine doses administered across the human population, as of February 3^rd^ 2023.^1^ Although approved COVID-19 vaccines have been effective in reducing mortality and morbidity caused by SARS-CoV-2 infection, the emergence of new variants that are able to evade the immune response has raised concerns about their long-term efficacy. Furthermore, the uneven distribution of vaccines worldwide has resulted in many low to middle income countries being left without access to variant-specific vaccines that are better suited for the evolving SARS-CoV-2 variant landscape. This highlights the need for the development of vaccines that can provide broad protection against a range of SARS-CoV-2 variants, as well as the importance of equitable distribution of vaccines to mitigate the risk of further virus evolution and spread.^2–5^ Since its emergence in late 2019, SARS-CoV-2 has continuously evolved, at a higher-than-expected rate, giving rise to multiple variants with multiple genetic mutations and various phenotypic properties, including increased transmissibility, virulence, and immune escape.^5,6^ The emergence of these variants has raised concerns about the efficacy of current vaccines and the potential for future outbreaks. Therefore, there is a critical need to develop effective vaccines that can provide broad and durable protection against SARS-CoV-2 and its variants. SARS-CoV-2 variants such as B.1.1.7 (Alpha), B.1.351 (Beta), and P.1 (Gamma) have exhibited substantial increases in immune escape from wildtype (WU) vaccine or infection induced immunity.^7,8^

The spike (S) protein of SARS-CoV-2 has been the main target of currently approved COVID-19 vaccines and of most COVID-19 vaccines in development.^9^ S protein allows for virus binding and infection of susceptible cells through interaction with host receptor angiotensin-converting enzyme 2 (ACE2).^10^ The S1 subunit of the S protein contains the receptor binding domain (RBD) that binds with ACE2, while the S2 subunit allows for cell fusion and viral entry.^11,12^ It has been widely acknowledged that antibodies targeting the S protein, particularly those binding to the RBD, are able to block the binding of SARS-CoV-2 to the cell receptor and prevent infection of susceptible cells.^13–17^ We have previously demonstrated the immunogenicity of S1 subunit targeting vaccines against various Beta-coronaviruses including SARS-CoV-1, SARS-CoV-2, and MERS.^18–23^

A focus for next-generation SARS-CoV-2 vaccine design is the investigation of novel vaccines which may be able to induce a broader immune response effective against multiple SARS-CoV-2 variants. A multivalent vaccine is a traditional approach used to increase antigen immunity coverage against multi-variant viruses such as SARS-CoV-2. We have previously demonstrated the immunogenicity of a trivalent protein subunit vaccine in BALB/c mice.^22^ Here, we assessed our S1 protein subunit vaccine, at an increased valency to tetravalent, in an advanced animal model more closely related to humans. Nonhuman primates (NHPs) are commonly used as preclinical models to evaluate the safety and efficacy of vaccines and therapeutics for infectious diseases, including SARS-CoV-2.^24–27^ We employed a rhesus macaque (RM) model of controlled simian immunodeficiency virus (SIV) infection to evaluate the immunogenicity of a tetravalent SARS-CoV-2 S1 protein subunit vaccine delivered with AddaVax adjuvant. Controlled SIV infection in RMs mimic a situation of chronic viral infection which can be encountered in humans, which may influence the development of immune responses to vaccination. Indeed, some studies reported lower SARS-CoV-2 antibody responses for people living with HIV.^28,29^ Several studies have demonstrated the utility of RMs as a preclinical model for SARS-CoV-2 vaccine development. For example, macaques have been used to evaluate the immunogenicity and the correlates of protection, as well as the protective efficacy of various vaccine platforms, including viral vector-based vaccines, mRNA vaccines, and protein subunit vaccines.^26,27, 30–34^ Moreover, the use of NHP models can provide critical insights into the mechanisms of vaccine-induced immunity, including the kinetics, specificity, and durability of the immune responses.

Here, we evaluated the immunogenicity of a tetravalent SARS-CoV-2 vaccine approach with S1 subunit protein vaccine targeting Wuhan S1, B.1.1.7 (Alpha), B.1.351 (Beta), and P.1 (Gamma). We chose these variants because, at the time of the start of the study, they represented a diverse and relevant set of SARS-CoV-2 strains that were circulating in different regions of the world and had distinct mutations in the spike protein, which is the main target of neutralizing antibodies. We found that vaccination induced robust humoral and cellular immune responses which resulted in antibodies capable of blocking ACE2 binding to 15 different SARS-CoV-2 variants, including multiple Omicron variants. Vaccination also induced antibodies that were able to block SARS-CoV-2 infection of susceptible cells by live wild-type (WU), Beta, and Delta variant viruses. We profiled the lymphocyte response to immunization for 2 months post initial prime vaccination through quantifying the number of T and B cells, investigating markers of T-cell activation, and memory subsets in peripheral blood mononuclear cells (PBMCs) and showed robust immune activation, primarily after boost immunization. We were also able to measure a spike-specific CD4^+^ T-cell response in the PBMC’s of RMs 42 days post-prime immunization, although, no CD8^+^ T-cell response was found. Our study further demonstrates the immunogenicity of protein subunit vaccines against SARS-CoV-2 targeting the S1 subunit of the spike protein while also contributing insights on approaches to further increase valency of currently approved COVID-19 vaccines.

## Results

### Design and expression or recombinant proteins

To produce recombinant proteins of SARS-CoV-2-S1 pAd/S1Wu, pAd/S1Alpha, pAd/S1Beta, and pAd/S1Gamma were generated by subcloning the codon-optimized SARS-CoV-2-S1 gene having C-tag into the shuttle vector, pAd (GenBank U62024) at *Sal* I and *Not* I sites (**Fig. 1A**). Variant-specific mutations for B.1.1.7 (Alpha), B.1.351 (Beta), and P.1 (Gamma) SARS-CoV-2 recombinant S1 proteins are outlined. To determine SARS-CoV-2-S1 expression from each plasmid, Expi293 cells were transfected with pAd/ S1WU, pAd/S1Alpha, pAd/S1Beta, and pAd/S1Gamma or pAd as a control. At 5 days after transfection, the supernatants of Expi293 cells were characterized by Western blot analysis. As shown in **Fig. 1B**, each S1 recombinant proteins were recognized by a polyclonal anti-spike of SARS-CoV-2 Wuhan antibody at the expected glycosylated monomeric molecular weights of about 110 kDa under the denaturing reduced conditions, while no expression was detected in the mock-transfected cells (lane1). The purified rS1WU, rS1Apha, rS1Beta, and rS1Gamma proteins using C-tagXL affinity matrix were determined by silver staining (**Fig. 1C**).

**Figure 1.**
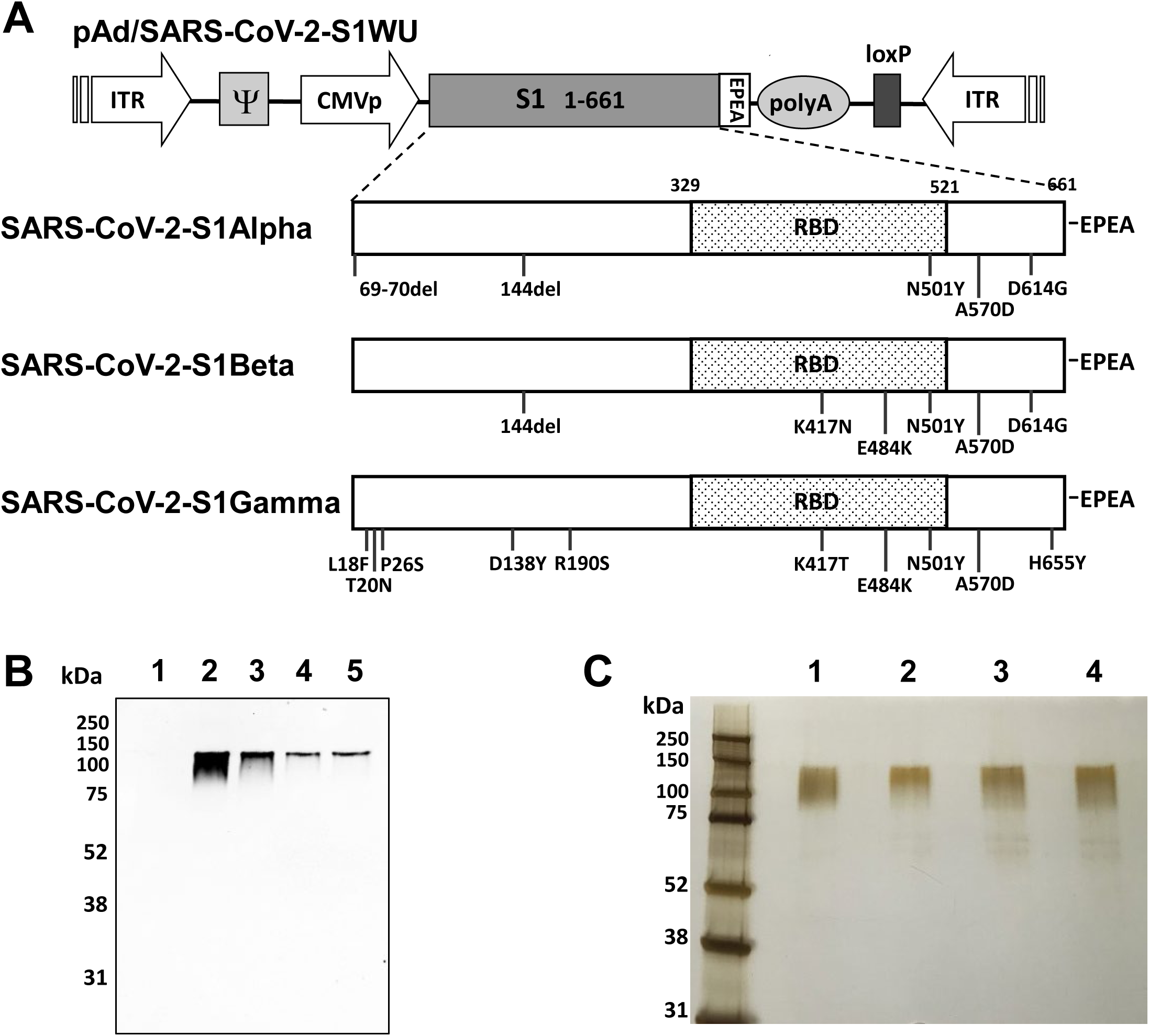
Construction and expression of tetravalent recombinant SARS-CoV-2-S1 proteins. **A.** A shuttle vector carrying the codon-optimized four variants of SARS-CoV-2-S1 gene encoding N-terminal 1-661 with c-tag (EPEA) was designated as shown in the diagram. Amino acid changes in the SARS-CoV-2-S1 region of in this study is shown. ITR: inverted terminal repeat; RBD: receptor binding domain. **B.** Detection of the SARS-CoV-2-S1 proteins by western blot with the supernatant of Expi293 cells transfected with pAd/S1WU (lane2), pAd/S1Alpha (lane3), pAd/S1Beta (lane4), and pAd/S1Gamma (lane5), respectively, using rabbit anti spike of SARS-CoV Wuhan polyclonal antibody. As a negative control, mock-transfected cells were treated the same (lane 1). **C.** Purified proteins, rS1WU (lane1), rS1Alpha (lane2), rS1Beta (lane3), and rS1Gamma (lane4), isolated by c-tag affinity purification were separated by SDS-PAGE and visualized by silver staining. Molecular weight marker (MW marker) is indicated on the left.

### Binding antibody and cross-variant live virus neutralizing antibody response

Prior to immunization, RMs were infected with a simian immunodeficiency virus (SIV) that naturally infects African green monkeys (SIVsab).^35^ This virus is completely controlled in RMs,^36^ in spite of retaining the replicative abilities.^37^ At the time of SARS-CoV-2 immunization, the RMs were controlling SIVsab for over a year. Upon prime and boost immunization, SIVsab viral loads remained undetectable suggesting no SIV activation upon vaccination. RMs were primed and boosted on week 3 with 60 µg total of rS1WU, rS1Apha, rS1Beta, and rS1Gamma, 15 µg of each antigen, mixed with 300 µl of AddaVax^TM^, squalene-based oil in water nano-emulsion adjuvant (**Fig. 2A**). To assess the magnitude of the antibody response we first determined Wuhan IgG antibody endpoint titers (EPT) in the sera of vaccinated RMs with ELISA. Serum samples collected prior to immunization, week 3, week 7, and week 9-11 after immunization were serially diluted to determine SARS-CoV-2-S1-specific IgG titers against Wuhan S1 using ELISA (**Fig. 2B**). RMs had detectable anti-S1 binding antibody response prior to immunization (**Fig 2B**), however, no neutralizing antibody response was found (**Fig. 2C**). S1-specific IgG titers were statistically increased at week 7 and week 9-11 when compared to week 0 (**Fig. 2B,** p < 0.05, Kruskal-Wallis test, followed by Dunn’s multiple comparisons). To evaluate the functional quality of vaccine-generate antigen-specific antibodies, we used a microneutralization assay (NT_90_) to test the ability of sera from immunized RMs to neutralize the infectivity of SARS-CoV-2. Sera, collected from RMs on week 3 (prior to booster immunization) and week 7 (4 weeks post boost) after primary immunization were tested for the presence of SARS-CoV-2-specific neutralizing antibodies with live SARS-CoV-2 Wuhan, Beta, and Delta viruses (**Fig. 2C**). High levels of neutralizing antibodies were detected in sera at week 3 and week 7 against Wuhan, Beta, and Delta SARS-CoV-2 variants (**Fig. 2C**) and showed a similar pattern with IgG endpoint titers in each RM (**Supplementary Fig. 2**). Furthermore, the geometric mean titers (GMT) of neutralizing antibodies at week 7 against the Wuhan, Beta, and Delta strain were increased with 6.4-, 5.4-, 3.2-fold compared at week 3, respectively, while only neutralizing antibody response against live Wuhan SARS-CoV-2 at week 7 was significantly increased when compared to preimmunized sera (**Fig. 2C**, p < 0.05, Kruskal-Wallis test, followed by Dunn’s multiple comparisons). Neutralization against highly immune-evasive Beta and Delta SARS-CoV-2 variants of concern (VOC) were found at slightly lower levels than Wuhan at both week 3 and week 7 (**Fig. 2C**). While Beta VOC S1 was included in the tetravalent immunization regimen, Delta VOC was not, highlighting the diverse response induced by tetravalent immunization in RMs.

**Figure 2.**
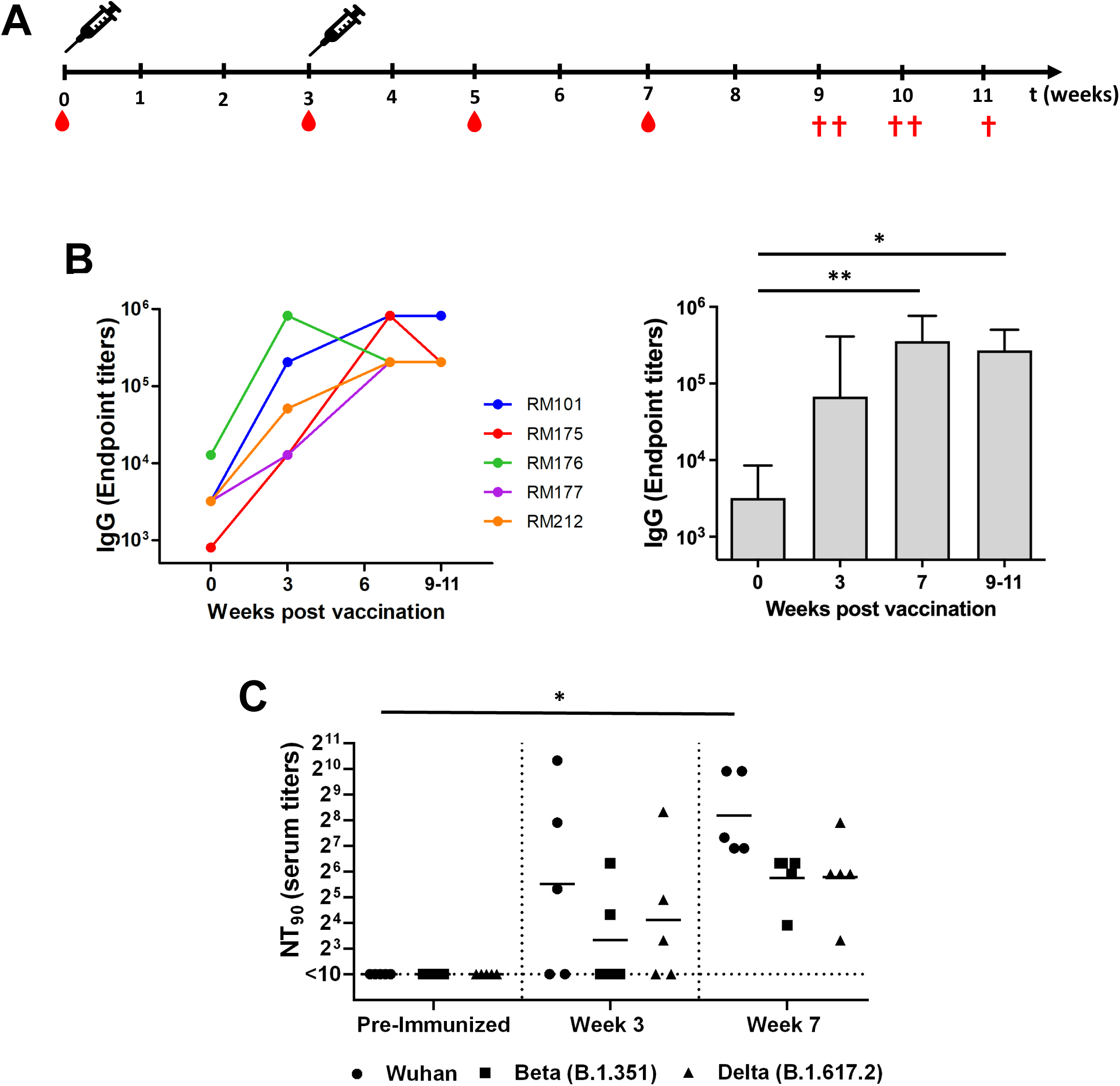
Antigen-specific antibody responses in rhesus macaques immunized with tetravalent SARS-CoV-2 rS1 protein subunit vaccine. **A.** Schedule of immunization and blood sampling for IgG end point titration. Rhesus macaques (N=5) were immunized with 60μg of tetravalent rS1 proteins of Wuhan, B.1.1.7 (Alpha), B.1.351 (Beta), and P.1 (Gamma) [15μg of each antigen] mixed with AddaVax adjuvant then administered to RMs arm at week 0 and 3. Syringes indicated the timing of immunization and the red drops denote times at which blood was drawn. The red crosses showed euthanized times of each RM. **B.** Sera were diluted and SARS-CoV-2-S1-specific antibodies were quantified by ELISA to determine the IgG endpoint titer. The IgG titers at each time points were showed in each RM. The bars represent geometric mean with geometric SD. **C.** Neutralizing antibodies in serum of mice prior to immunization, along with week 3 and week 7 post immunization were measured using a microneutralization assay (NT_90_) with SARS-CoV-2 Wuhan, Beta, and Delta. Serum titers that resulted in 90% reduction in cytopathic effect compared to the virus control were reported. Horizontal lines represent geometric mean titers. Groups were compared by Kruskal-Wallis test at each time point, followed by Dunn’s multiple comparisons. Significant differences are indicated by *p < 0.05. N = 5 rhesus macaques per group for each experiment.

### Potent ACE2 binding inhibition effective against 15 different SARS-COV-2 VOC’s spikes

For further insight into the neutralizing capabilities of antibodies induced by vaccination we used the Meso Scale Discovery (MSD) V-PLEX SARS-CoV-2 (ACE2) Kit to measure the inhibition of binding between angiotensin converting enzyme-2 (ACE2) and trimeric spike protein of SARS CoV-2 variants. Initially, we used kit Panel 18 including Wuhan S and spikes from variants; Alpha (B.1.1.7), Beta (B.1.351), Gamma (P.1), Delta (B.1.617, B.1.617.2), Zeta (P.2), Kappa (B.1.617.1), B.1.526.1, B.1.617, and B.1.617.3 (**Fig. 3**). Sera from vaccinated RMs were examined at week 7, due to that being the peak of measured IgG binding antibody response and compared to preimmunized sera (**Fig. 2A, Fig. 3**). Antibodies blocking ACE2 and trimeric S binding of all variants, by over 90% inhibition, were detected in all 1:10 diluted RM sera at Week 7 (**Fig. 3**). Week 7 sera ACE2 binding inhibition for RMs was significantly increased, when compared to preimmunized sera, for Wuhan, B.1.1.7, B.1.351, P.1, B.1.617.2, P.2, B.1.617.1, B.1.526.1, B.1.617, and B.1.617.3 Spike (**Fig. 3**, p < 0.05, Mann-Whitney Test).

**Figure 3.**
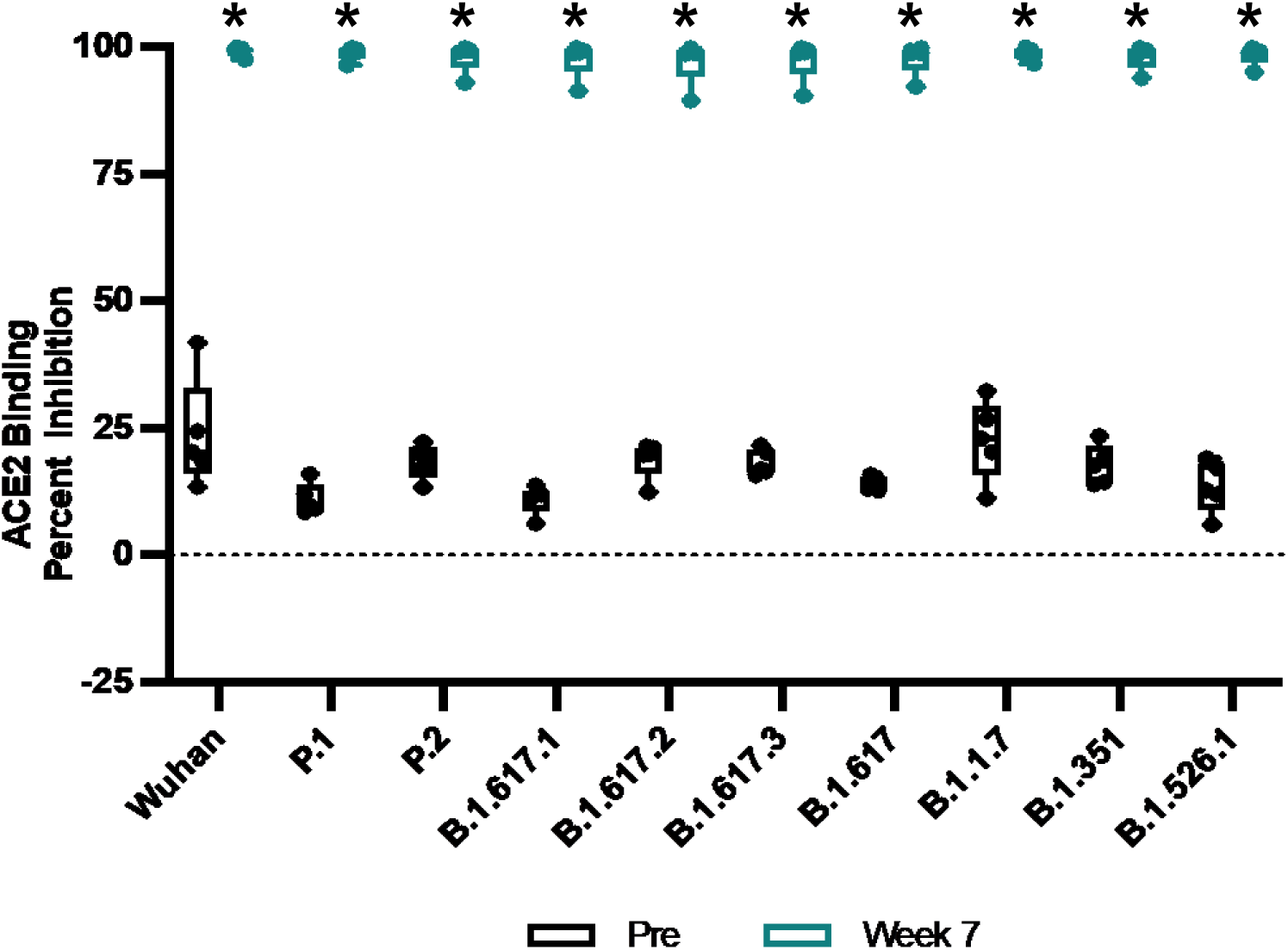
Percent ACE2 binding inhibition of neutralizing antibodies against SARS-CoV-2 variants. Antibodies in sera (diluted 1:10) capable of neutralizing the interaction between SARS-CoV-2 Wuhan, Alpha (B.1.1.7), Beta (B.1.351), Gamma (P.1), Delta (B.1.617.2), Zeta (P.2), Kappa (B.1.617.1), New York (B.1.516.1), India (B.1.617 and B.1.617.3) variants spike and ACE2 were examined in all animals preimmunization and Week 7 post prime immunization with V-PLEX SARS-CoV-2 Panel 18. Groups were compared by Kruskal-Wallis test at each time point, to preimmunized sera control, followed by Dunn’s multiple comparisons. Significant differences are indicated by *p < 0.05. N = 5 rhesus macaques per group for each experiment.

To assess the neutralizing capabilities of RM vaccine induced antibodies against Omicron (BA.1) VOC, and Omicron sub-variants (BA.2, BA.3, BA.1+R346K, BA.1+L452R) we used MSD V-Plex SARS-CoV-2 ACE2 Kit Panel 25 (**Fig. 4**). Panel 25 includes SARS-CoV-2 Wuhan, BA.1, BA.2, AY.4, BA.3, BA.1+R346K, BA.1+L452, B.1.1.7, B.1.351, and B.1.640.2 trimeric spike. Sera from vaccinated RMs were examined at week 3, week 7, and week 9-11 post vaccination and compared to preimmunized sera at a 1:10 dilution (**Fig. 4A**) and 1:100 dilution (**Fig 4B**). Week 7 and Week 9-11 RM sera ACE2-binding inhibition were significantly increased when compared to preimmunized sera for Wuhan, AY.4 (Delta lineage), BA.1+L452R, B.1.1.7, B.1.351, and B.1.640.2 VOC spikes at 1:10 dilution (**Fig. 4A**, p < 0.05, Kruskal-Wallis test, followed by Dunn’s multiple comparisons). Week 7 RM sera ACE2-binding inhibition were significantly increased when compared to preimmunized sera for BA.1 VOC spike at 1:10 dilution (**Fig. 4A** p < 0.05, Kruskal-Wallis test, followed by Dunn’s multiple comparisons).While not statistically significantly increased when compared to preimmunized RM sera; RMs demonstrated moderate ACE2-binding inhibition for BA.2, BA.3, and BA.1+R346K VOC spikes weeks 7 and 9-11 post immunization at 1:10 dilution (**Fig. 4A**, p > 0.05, Kruskal-Wallis test, followed by Dunn’s multiple comparisons). To further interrogate the vaccine-induced neutralizing capabilities of RMs, we further substantially diluted RM sera to 1:100 (**Fig. 4B**). Week 7 RM 1:100 diluted sera ACE-2 binding inhibition was significantly increased when compared to preimmunized sera for Wuhan, AY.4, B.1.1.7, B.1.351, B.1.640.2 VOC spikes (**Fig. 4B**, p < 0.05, Kruskal-Wallis test, followed by Dunn’s multiple comparisons). At 1:100 dilution, RM sera did not have ACE-2 binding inhibition above preimmunized sera for BA.1, BA.2, BA.3, BA.1+R346K, BA.1+L452R VOC spikes (**Fig. 4B**). Results suggest the necessity of the booster immunization to induce potent and cross variant recognizing antibodies. Results also suggest that vaccination induced antibodies that are able to potently recognize and block ACE2 binding of a wide range of SARS-CoV-2 variants spikes by week 7 post prime immunization.

**Figure 4.**
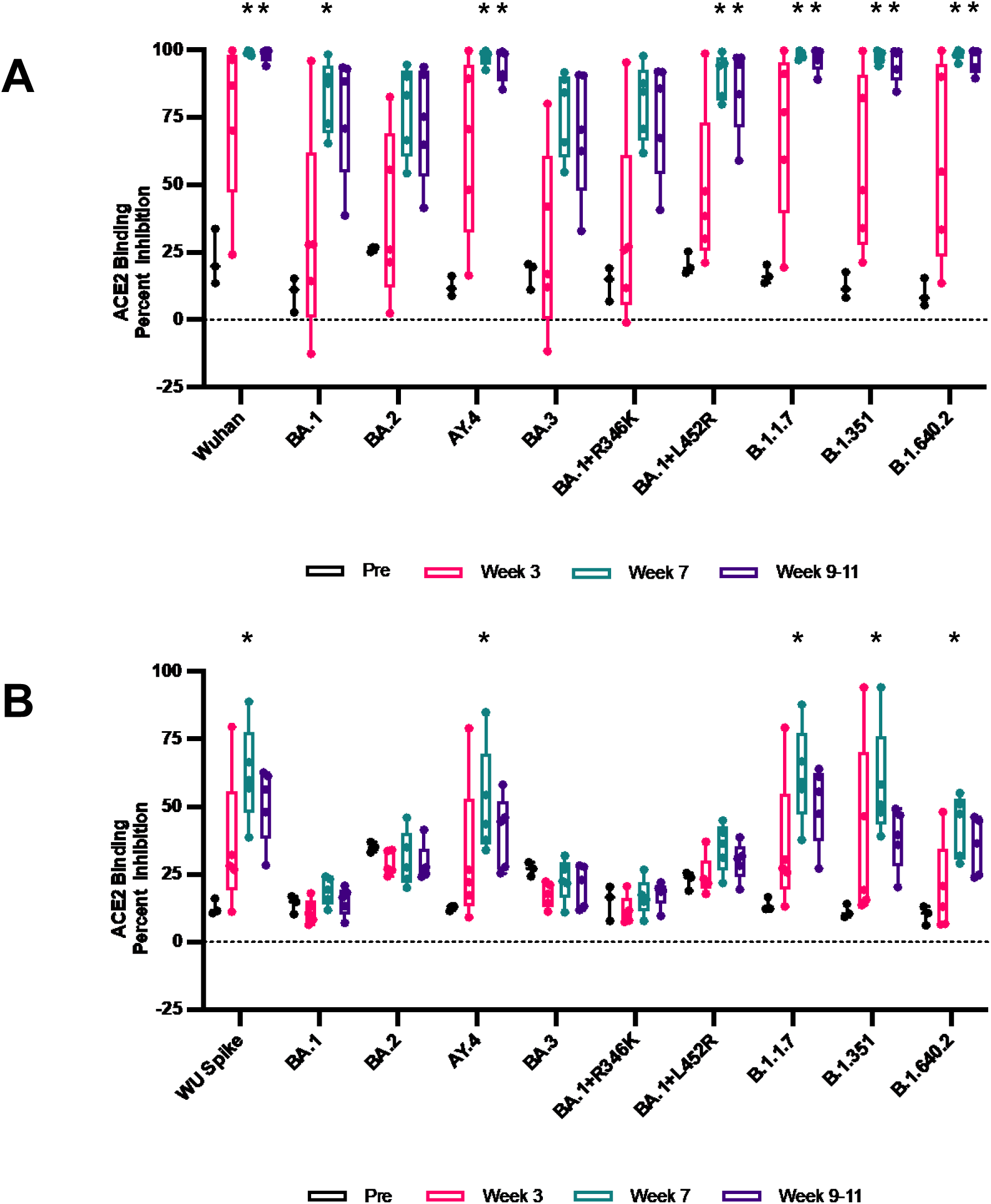
Percent ACE2 binding inhibition of neutralizing antibodies against Omicron SARS-CoV-2 variants. Antibodies in sera, diluted **A.** 1:10 and **B.** 1:100 capable of blocking the binding of SARS-CoV-2 spike including Wuhan and spikes from immune evasive variants; BA.1, BA.2, AY.4 (Delta lineage), BA.3, BA.1+R346K mutation, BA.1+L452R mutation, B.1.1.7 (Alpha), B.1.351 (Beta), and B.1.1640.2 to ACE2 were detected with a V-PLEX SARS-CoV-2 Panel 25. Groups were compared by Kruskal-Wallis test at each time point, to preimmunized sera control, followed by Dunn’s multiple comparisons. Significant differences are indicated by *p < 0.05. N = 5 rhesus macaques per group for each experiment.

### Longitudinal lymphocyte dynamics and cell-mediate immune response to vaccination shows immune activation primarily observed after boost

To investigate the kinetics and magnitude of immune responses induced by the tetravalent SARS-CoV-2 vaccine, we monitored the peripheral blood mononuclear cells (PBMCs) of vaccinated rhesus macaques over a 60-day period. PBMCs are a mixture of different immune cell types, including T cells and B cells, and are a useful tool for investigating the immune response to vaccination in vivo.

**Fig. 5** shows the dynamics of CD3^+^ T-cells (**Fig. 5A**), CD4^+^ T-cells (**Fig. 5B**), CD8^+^ T-cells (**Fig. 5C**), and CD20^+^ B cell (**Fig. 5D**) counts over 60 days. We observed increases in all T-cell subsets (CD3^+^, CD4^+^, and CD8^+^) and B cells (CD20^+^) after the prime and especially after the boost, demonstrating clear increases for all subsets, with the CD8^+^ T cell count showing the greatest increase after boost immunization compared to the other cell types.

**Figure 5.**
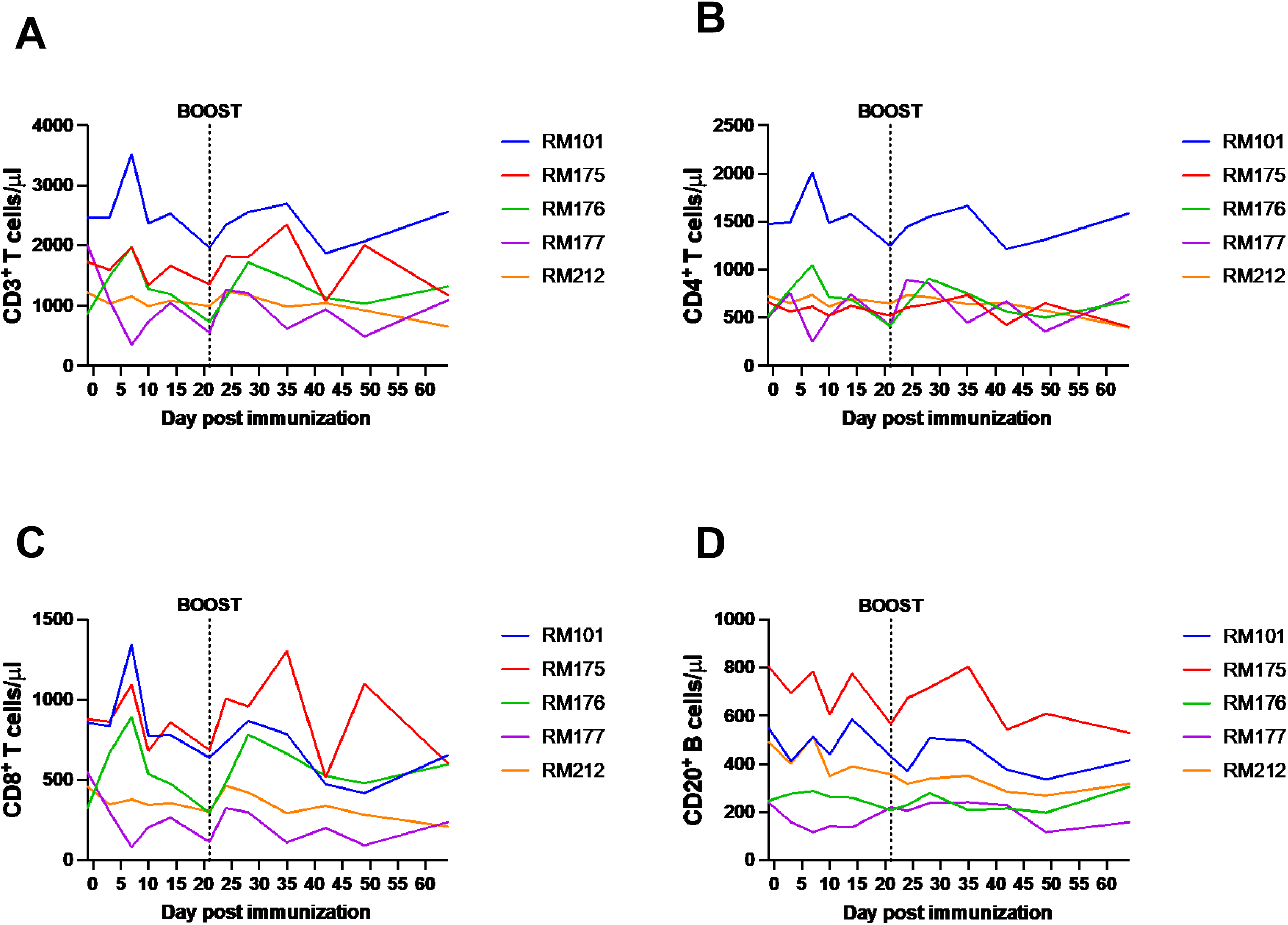
CD3, CD4, CD8, and CD20 cell counts post immunization and boost. Absolute counts of immune cells in whole blood and immunophenotyping of circulating immune cells were determined by flow cytometry. 50 μl of whole blood were added to a TruCount tube (BD Biosciences) containing an antibody mix, allowing to precisely quantify **A.** CD45^+^ cells, **B.** CD4^+^, **C.** CD8^+^ T cells, and **D.** CD20^+^ B cells in blood per μl. PMBC’s from RMs were collected and analyzed on Days -1, 3, 7, 10, 14, 21, 24, 28, 31, 35, 42, 49, and 64 days post prime immunization. Individual results for each RM are depicted.

**Fig. 6** shows the fraction of activating and proliferating CD4^+^ and CD8^+^ T cells. We used the activation markers CD69 and HLDR and CD38, as previously described in the literature.^38–40^ We also used Ki-67 as a marker for cell proliferation. CD69^+^ CD4^+^ T-cell induction was mainly observed in RM177 (**Fig. 6A**). Ki67^+^ CD4^+^ T cells showed moderate increases in percentage after boost vaccination (**Fig. 6B**). HLA-DR^+^ CD38^+^ CD4^+^ T-cells showed activation post prime and boost with a return to near baseline by Day 40 (**Fig. 6C**). The fraction of CD69^+^ CD8^+^ T-cells increased in all RMs post prime and boost, with most starting to return to prevaccination levels at day 60 (**Fig. 6D**). The induction of Ki-67^+^ CD8^+^ T-cells was primarily seen at day 40 postimmunization (**Fig. 6E**), while HLA-DR^+^ CD38^+^ CD8^+^ T-cell activation was mainly seen in RM175 and RM176 at different timepoints (**Fig. 6F**). However, the induction of HLA-DR^+^ CD38^+^ CD8^+^ T cells was not as robust as that of CD69^+^ CD8^+^ T cells and Ki-67^+^ CD8^+^ T cells (**Fig. 6F, Fig. 6D, Fig. 6E**).

**Figure 6.**
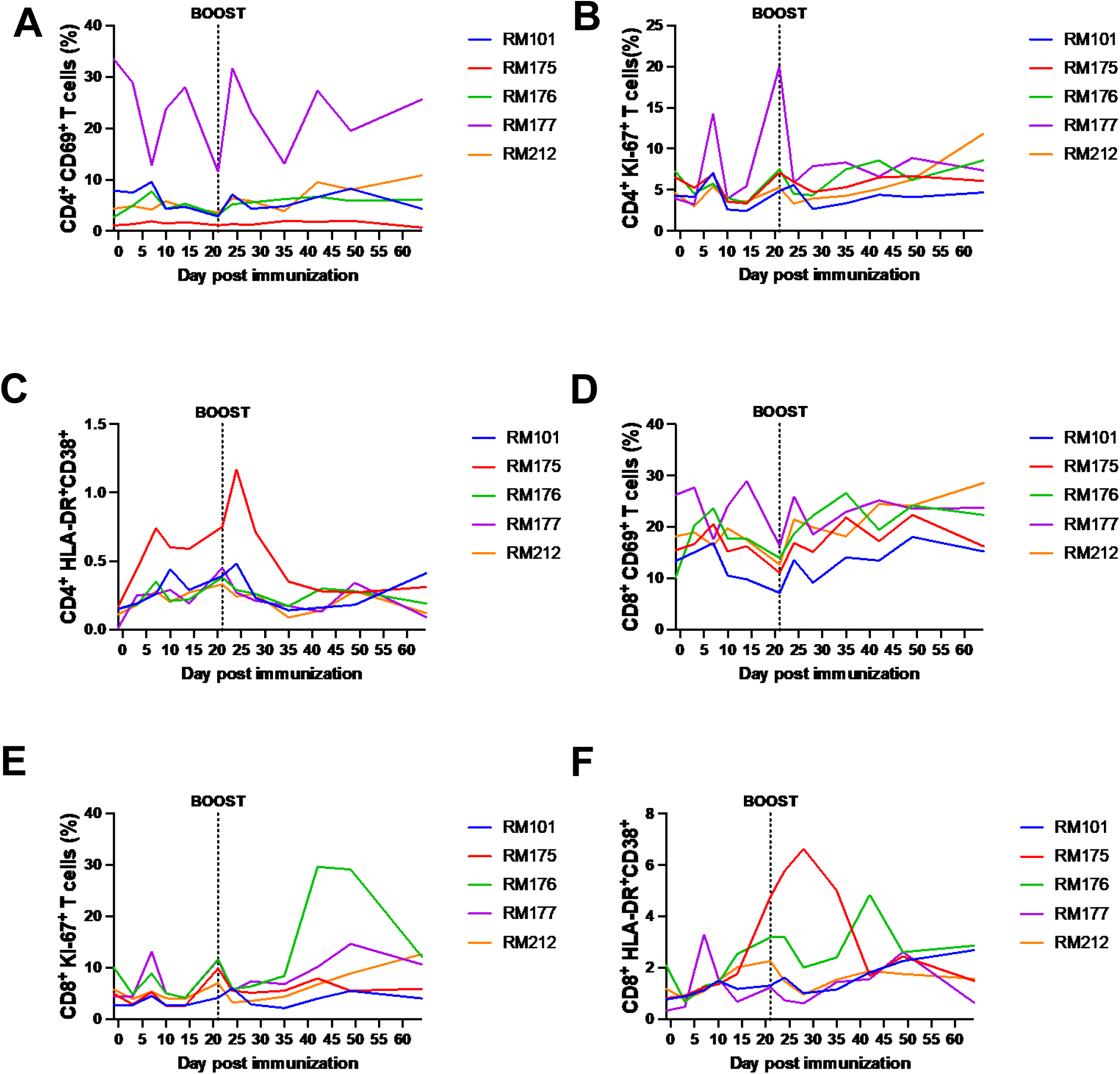
CD4 CD8 T cell activation post immunization and boost. Whole peripheral blood was stained with fluorescently labeled antibodies for CD4^+^, CD8^+^, CD69^+^, Ki-67^+^, and HLA-DR^+^ to investigate CD4 and CD8 activation induced by vaccination with flow cytometry. **A.** Frequencies of CD4^+^ CD69^+^ T cells, **B.** Frequencies of CD4^+^ Ki-67^+^ T cells, **C.** Frequencies of CD4^+^ HLA-DR^+^ CD38^+^ T cells, **D.** Frequencies of CD8^+^ CD69^+^ T cells, **E.** Frequencies of CD8^+^ Ki-67^+^ T cells, and **F.** Frequencies of CD8^+^ HLA-DR^+^ CD38^+^ T cells. PMBC’s from RMs were collected and analyzed on Days -1, 3, 7, 10, 14, 21, 24, 28, 31, 35, 42, 49, and 64 days post prime immunization. Individual results for each RM are depicted.

**Fig. 7** shows the changes in the distribution of T-cell memory subsets over time. We defined naïve, central memory (CM), and effector memory (EM) T cells using CD28^+^ and CD95^+^ markers. Naïve T cells are CD28^+^ CD95^neg^, CM T-cells are CD28^+^ CD95^+^, and EM T cells are CD28^neg^ CD95^+^. We observed that both CD4^+^ and CD8^+^ central memory T cells (**Fig. 7A & 7D**), along with naïve CD4^+^ naïve CD8^+^ T cells (**Fig. 7C & 7F**), decreased in abundance after prime and boost, while CD4^+^ and CD8^+^ effector memory T cells (**Fig. 7B & 7E**) increased in abundance after prime boost. This finding suggests that the tetravalent S1 protein vaccine induces a shift towards an effector memory phenotype and away from a central memory phenotype, which may be beneficial in generating a rapid and robust response to vaccination.

**Figure 7.**
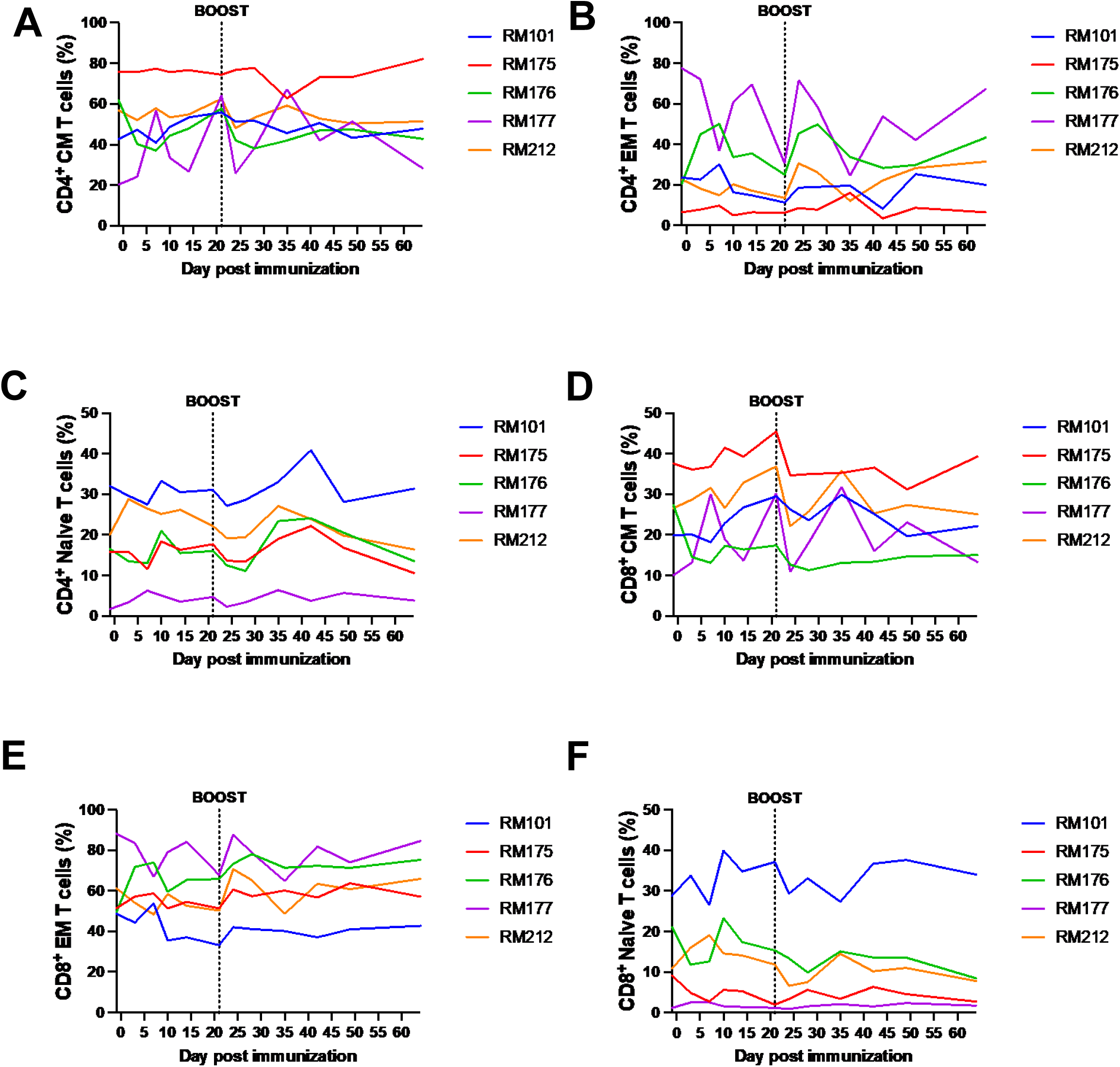
T cell memory subset dynamics and induction post immunization and boost. Whole peripheral blood was stained with fluorescently labeled antibodies for CD4^+^, CD8^+^, CD28^+^ and CD95^+^. Memory subsets were defined naive, central memory (CM), and effector memory (EM) T cells using CD28^+^ and CD95^+^ markers. Naive T cells are CD28^+^CD95-, CM T cells are CD28^+^CD95^+^, and EM T cells are CD28-CD95^+^. **A.** Frequencies of CD4^+^ CM T cells, **B.** Frequencies of CD4^+^ EM T cells, **C.** Frequencies of CD4^+^ Naive T cells, **D.** Frequencies of CD8^+^ CM T cells, **E.** Frequencies of CD8^+^ EM T cells, and **F.** Frequencies of CD8^+^ Naïve T cells. PMBC’s from RMs were collected and analyzed on Days -1, 3, 7, 10, 14, 21, 24, 28, 31, 35, 42, 49, and 64 days post prime immunization. Individual results for each RM are depicted.

Intracellular cytokine staining was performed to evaluate the spike-specific T-cell responses in CD4^+^ and CD8^+^ T cells after stimulation with a spike peptide pool at day 0 and day 42 postvaccination in PBMCs (**Fig. 8**). We tested for interferon-gamma (IFN-γ), interleukin-2 (IL-2), and tumor necrosis factor-alpha (TNF-α) cytokine staining. Only RM212 induced an IFN-γ CD4^+^ T-cell response, while no such response was observed in the other four RMs (**Fig. 8A**). In **Fig. 8B**, we observed an induction of IL-2 CD4^+^ T-cell response in RM212 and to a lesser extent in RM101, but not in the other three RMs. **Fig. 8C** shows an induction of TNFα CD4^+^ T-cell response in RM212, RM176 and, to a minimal extent, in RM101, RM175, and RM177. Notably, we were not able to detect a spike specific CD8^+^ T-cell response at day 0 or day 42 post vaccination (data not shown). RM212 mounted a robust CD4^+^ T-cell response for all three cytokines at day 42. These results suggest that there is a variable induction of cytokine responses in CD4^+^ T cells among different RMs at day 42 postvaccination.

**Figure 8.**
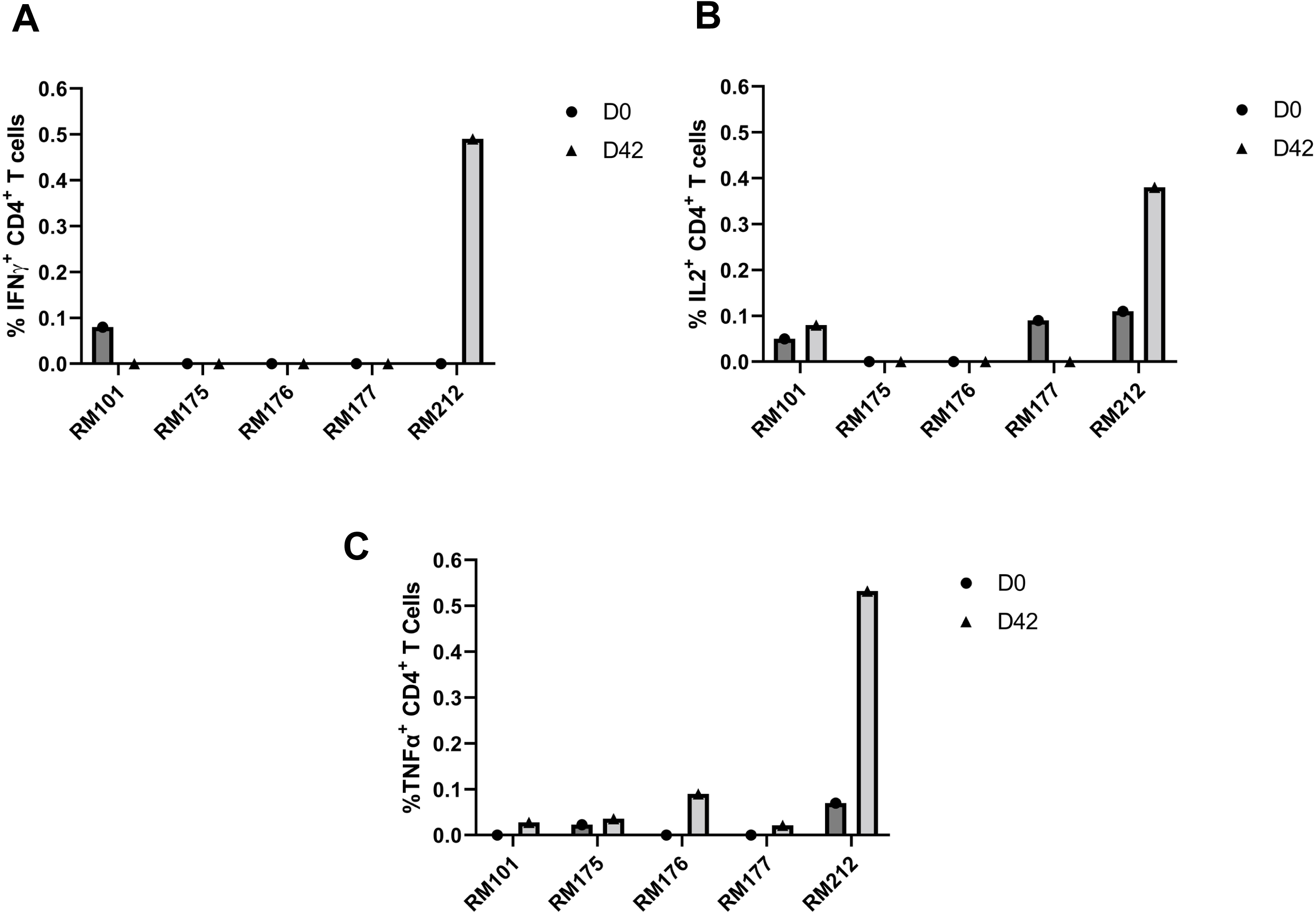
Spike-specific CD4+ T cell responses at Day 0 and Day 42 post immunization in PBMC’s. PBMC’s collected prior to immunization and on Day 42 post prime immunization were stimulated with PepTivator SARS-CoV-2-S1 (a pool of S1 MHC class I– and MHC class II– restricted peptides), followed by intracellular staining (ICS) and flow cytometry to identify SARS-CoV-2 S1 specific T cells. **(A)** Frequencies of SARSLCoVL2 S1 CD4^+^ IFNLγ^+^ T cells. Individual results for each RM are depicted. **(B)** Frequencies of SARSLCoVL2 S1 CD4^+^ IL-2^+^ T cells. Individual results for each RM are depicted. **(C)** Frequencies of SARSLCoVL2 S1 CD4^+^ TNFα T cells. Individual results for each RM are depicted. Day 0 PBMC responses are indicated by solid circle. Day 42 PBMC responses are indicated by solid triangle.

Overall, the use of PBMC’s allowed for the unique assessment of the dynamics of immune activation after vaccination. The results showed a clear increase in T-cell counts and activation after boost immunization, with the CD8^+^ T-cell counts showing the greatest increase. The use of CD markers allowed for the differentiation of T-cell subsets and their activation status, with the CD8^+^ T cells expressing either CD69 or Ki-67 CD8^+^ T cells showing the most robust dynamics. Additionally, there was evidence of a functional spike-specific CD4^+^ T-cell response in RMs at day 42 post vaccination, albeit in the context of no CD8^+^ T-cell response. These findings highlight the potential of this vaccine candidate to induce a robust cellular immune response, which is critical for controlling viral infections.

## Discussion

We evaluated the immunogenicity and efficacy of a tetravalent COVID-19 vaccine candidate based on the spike S1 protein of SARS-CoV-2 in an NHP model of controlled SIV infection. RMs infected with SIVsab from African green monkeys are able to control viral replication and disease progression through maintaining a healthy immune system, unlike HIV-1 in humans.^36^ The SIVsab-infected RMs in this study were elite controllers for about a year prior to SARS-CoV-2 immunization.

There were weaker band in western blot of the supernatant after a transient transfection with pAd/S1Alpha, pAd/S1Beta, and pAd/S1Gamma compared with pAd/S1WU **(Fig. 1B)**, which might be explained by the usage of anti-spike of SARS-CoV-2 Wuhan as a primary antibody. Indeed, no big differences were observed in yield pre or post C-tag purification of each recombinant proteins after transient transfection by sandwich ELISA with standard of each purified rS1 proteins **(Supplementary Fig.1).**

Our vaccine formulation induced high levels of binding antibodies against the Wuhan strain of SARS-CoV-2, as well as neutralizing antibodies against live B.1.351 (Beta), and B.1.617.2 (Delta) VOC (**Fig. 2**). The sera of vaccinated RMs exhibited potent ACE2-binding inhibition capabilities against a suite of SARS-CoV-2 VOC spikes including Omicron (BA.1) and Omicron subvariants (BA.2, BA.3, BA.1+R246K, and BA.1+L452R) (**Fig. 3 & Fig. 4**). These findings are consistent with previous studies demonstrating the immunogenicity and cross-reactivity of COVID-19 vaccines NHP models.^26,27, 30–33,41^

Importantly, the vaccine candidate also induced cellular immune responses, including T cell responses, which have been shown to play a critical role in COVID-19 immunity and protection.^42–49^ We investigated the cellular immune response to the tetravalent SARS-CoV-2 vaccine in vaccinated RMs, using a range of markers to examine T-cell subsets and activation status. The results showed that all T-cell subsets and B cells increased after the prime and especially after the boost, with the CD8^+^ T-cell count showing the greatest increase after boost immunization compared to other cell types (**Fig. 5**). We demonstrate that the tetravalent S1 subunit protein COVID-19 vaccine candidate induces CD4^+^ and CD8^+^ T-cell activation, as indicated by increased expression of CD69, HLA-DR, CD38, and Ki-67 activation and proliferation markers on both T-cell subsets (**Fig. 6**). The distribution of T-cell memory subsets over time was also investigated, revealing a decrease in abundance of both CD4^+^ and CD8^+^ central memory T cells, along with CD4^+^ and CD8^+^ naive T cells after prime and boost (**Fig. 7**). In contrast, CD4^+^ and CD8^+^ effector memory T cells increased in abundance after prime boost, indicating a shift towards an effector memory phenotype and away from a central memory phenotype induced by the tetravalent S1 protein vaccine (**Fig. 7**). Furthermore, intracellular cytokine staining was performed to evaluate the spike-specific responses of CD4^+^ and CD8^+^ T cells after stimulation with a spike peptide pool (**Fig. 8**). Cytokine staining for IFN-γ, IL-2, and TNF-α was tested and a variable induction of cytokine responses by CD4^+^ T cells among different RMs at day 42 postvaccination was observed (**Fig. 8**). However, no spike-specific response of the CD8^+^ T cells was detected at day 0 or day 42. It is possible that the spike-specific CD8^+^ T cells were present, but were not detected by the intracellular staining assay, as this assay may not be sensitive enough to detect low-frequency antigen-specific CD8^+^ T cells. It is also possible that the undetectable spike-specific CD8^+^ T-cell response at day 42 post-vaccination was related to the time-point used, which was too late after boost, such as the vaccine-specific T cells had already started to wane in abundance, as shown by Arunachalam et al.^50^ Altogether, our study demonstrates that the tetravalent S1 protein vaccine candidate was able to induce a robust SARS-CoV-2-specific immune response in RMs, which is promising for future development and testing of COVID-19 vaccines in humans.

The results of our study have important implications for COVID-19 vaccine development and implementation in humans. The vaccine candidate induced not only humoral immune responses but also cellular immune responses, which have been shown to be important for long-term immunity.^51^ The use of RMs as an animal model for studying vaccine efficacy has been widely accepted in the scientific community.^25,26,34,52^ Here we have used RM controllers based on the rationale that SIV controllers have a nearly healthy immune system (able to control SIV replication).^36^ We also wanted to assess whether the induction of T-cell activation at the effector sites would result in a burst of SIV replication. Such a boosting of SIV was reported to occur after administration of vectorized vaccines.^53^ The use of NHP models has been shown to be highly informative for predicting vaccine efficacy in humans.^54,55^

The results showed that the vaccine induced both humoral and cellular immune responses against SARS-CoV-2, including neutralizing antibodies, ACE2 blocking antibodies, and T-cell responses. Furthermore, the vaccine candidate was able to generate Omicron variant binding and ACE2 blocking antibodies without specifically vaccinating with Omicron, suggesting the potential for broad protection against emerging variants.^56–60^ This is particularly significant given the emergence of highly diverged SARS-CoV-2 variants, such as Omicron, which have raised concerns about vaccine efficacy and the need for updated vaccines.^56,58,59,61^ Another significant feature of the vaccine candidate is its tetravalent composition, which targets the spike proteins of four different SARS-CoV-2 variants. This approach has the potential to provide broad protection against multiple SARS-CoV-2 variants, as well as to minimize the risk of immune escape and emergence of new variants.

Protein subunit vaccines are known for their safety, ease of large-scale production, and distribution, and have been used in other successful vaccine campaigns, such as the hepatitis B vaccine.^54,62–64^ This makes protein subunit vaccines an ideal candidate for worldwide vaccine equity, particularly for countries that may not have access to the more complex mRNA or viral vector vaccine platforms. Furthermore, the ability to store and transport protein subunit vaccines at a relatively low temperature (−20°C to 4°C), compared to the ultra-low temperature required for mRNA vaccines, makes their distribution and administration easier in resource-limited settings.^65,66^ The protein subunit platform is also amenable to alternative routes of administration, such as intradermal delivery, which has been shown to increase immunogenicity in other vaccine studies.^20,67–69^ In summary, the tetravalent S1 protein subunit vaccine represents a promising vaccine candidate against SARS-CoV-2, particularly for populations that may not have access to other vaccine platforms and could potentially be further optimized to enhance its immunogenicity.

However, it should be noted that this study has limitations. The sample size was small and we did not perform a SARS-CoV-2 virus challenge in our vaccinated RMs to fully assess vaccine efficacy.^27,50^ While our results show promising immune responses to the tetravalent SARS-CoV-2 vaccine in RMs, a virus challenge would have provided further insights into the effectiveness of the vaccine in preventing infection and disease. Additionally, our study did not evaluate the durability of the antibody response generated by the vaccine over a longer period. Studies have shown that antibody responses to SARS-CoV-2 vaccines may wane over time, which highlights the importance of evaluating the longevity of vaccine-induced immunity.^70–75^ Finally, we did not assess mucosal immunity in our study, which is an important aspect of immune protection against respiratory viruses like SARS-CoV-2. Mucosal immunity may provide an additional layer of protection against infection and transmission, and future studies should investigate the mucosal immune response to the tetravalent SARS-CoV-2 vaccine.^31,76–79^

The tetravalent S1 subunit protein COVID-19 vaccine candidate evaluated in this study contained SARS-CoV-2 S1 antigens from the Wuhan strain, as well as the B.1.1.7 variant, B.1.351 variant, and P.1 variant. Our study demonstrates that this vaccine candidate can induce both humoral and cellular immune responses, as evidenced by increased cell counts in both T and B cells, and the production of neutralizing and cross-reactive antibodies, as well as ACE2 blocking antibodies and T cell responses. It is important to note that the RMs used in this study were infected with SIVsab and controlled the infection for a year prior to immunization. The ability of these animals to control the SIVsab infection, without reactivation of virus upon immunization, while mounting immune responses to the vaccine candidate, further demonstrates the potential of this vaccine candidate to provide robust protection against SARS-CoV-2, even in individuals with pre-existing conditions. Moreover, the tetravalent composition of the vaccine candidate has significant implications for COVID-19 vaccine development and implementation, with the potential to provide broad protection against multiple SARS-CoV-2 variants and to minimize the risk of immune escape and emergence of new variants.

## Materials and methods

### Construction of recombinant protein expressing vectors

The coding sequence for SARS-CoV-2-S1 amino acids 1 to 661 of full-length from BetaCoV/Wuhan/IPBCAMS-WH-05/2020 (GISAID accession id. EPI_ISL_ 403928) having C-terminal tag known as ‘C-tag’, composed of the four amino acids (aa), glutamic acid-proline-glutamic acid-alanine (E-P-E-A) flanked with *Sal* I & *Not* I was codon-optimized using the UpGene algorithm for optimal expression in mammalian cells (68) and synthesized (GenScript). The construct also contained a Kozak sequence (GCCACC) at the 5′ end. For Alpha variant (B.1.1.7), SARS-CoV-2-S1 mutated Del69-70; Del144; N501Y; A570D; D614G was synthesized. Also, Beta variant (B.1.351) of SARS-CoV-2-S1 (Del144; K417N; E484K; N501Y; A570D; D614G) and Gamma variant (P.1) of SARS-CoV-2-S1 (L18F; T20N; P26S; D138Y; R190S; K417T; E484K; N501Y; H655Y) were synthesized based on above codon-optimized SARS-CoV-2-S1 Wuhan. pAd/S1WU, pAd/S1Alpha, pAd/S1Beta, and pAd/S1Gamma, were then created by subcloning the four variants of codon-optimized SARS-CoV-2-S1 inserts into the shuttle vector, pAdlox (GenBank U62024), at *Sal* I/*Not* I sites. The plasmid constructs were confirmed by DNA sequencing.

### Transient Production in Expi293 Cells

pAd/S1WU, pAd/S1Alpha, pAd/S1Beta, and pAd/S1Gamma, were amplified, and purified using ZymoPURE II plasmid maxiprep kit (Zymo Research). For Expi293 cell transfection, we used ExpiFectamie^TM^ 293 Transfection Kit (ThermoFisher) and followed the manufacturer’s instructions. Cells were seeded 3.0 × 10^6^ cells/ml one day before transfection and grown to 4.5-5.5 × 10^6^ cells/ml. 1μg of DNA and ExpiFectamine mixtures per 1ml culture were combined and incubated for 15 min before adding into 3.0L×L10^6^ cells/ml culture. At 20 h post-transfection, enhancer mixture was added, and culture was shifted to 32°C. The supernatants were harvested 5 days post transfection and clarified by centrifugation to remove cells, filtration through 0.8 μm, 0.45 μm, and 0.22 μm filters and either subjected to further purification or stored at 4°C before purification.

### SDS-PAGE and western blot

To evaluate the expression of S1 from the plasmids, Expi293 cells were transfected with pAd/S1WU, pAd/S1Alpha, pAd/S1Beta, and pAd/S1Gamma, respectively. At 5 days after transfection, 10 μl each supernatant of Expi293 cells was subjected to sodium dodecyl sulfate polyacrylamide gel electrophoresis (SDS-PAGE) and Western blot as previously described.^20^ Briefly, after the supernatants were boiled in Laemmli sample buffer containing 2% SDS with beta-mercaptoethanol (β-ME), the proteins were separated by Tris-Glycine SDS-PAGE gels and transferred to nitrocellulose membrane. After blocking for 1 hour at room temperature (RT) with 5% non-fat milk in PBST, rabbit anti-SARS-CoV Wuhan spike polyclonal antibody (1:3000) (Sino Biological) was added and incubated overnight at 4 °C as primary antibody, and horseradish peroxidase (HRP)-conjugated goat anti-rabbit IgG (1:10000) (Jackson immunoresearch) was added and incubated at RT for 2 hours as secondary antibody. After washing three times with PBST, the signals were visualized on an iBright FL 1500 Imager (ThermoFisher).

### Purification of recombinant proteins

The recombinant proteins named rS1WU, rS1Alpha, rS1Beta, and rS1Gamma were purified using a CaptureSelect^TM^ C-tagXL Affinity Matrix prepacked column (ThermoFisher) and followed the manufacturer’s guidelines. Briefly, The C-tagXL column was conditioned with 10 column volumes (CV) of equilibrate/wash buffer (20 mM Tris, pH 7.4) before sample application. Supernatant was adjusted to 20 mM Tris with 200 mM Tris (pH 7.4) before being loaded onto a 5-mL prepacked column per the manufacturer’s instructions at 5 ml/min rate. The column was then washed by alternating with 10 CV of equilibrate/wash buffer, 10 CV of strong wash buffer (20 mM Tris, 1 M NaCl, 0.05% Tween-20, pH 7.4), and 5 CV of equilibrate/wash buffer. The recombinant proteins were eluted from the column by using elution buffer (20 mM Tris, 2 M MgCl_2_, pH 7.4). The eluted solution was concentrated and desalted with preservative buffer (PBS) in an Amicon Ultra centrifugal filter devices with a 50,000 molecular weight cutoff (Millipore). The concentrations of the purified recombinant proteins were determined by the BCA protein assay kit (ThermoFisher) and separated by reducing SDS-PAGE and visualized by silver staining. The rest proteins were aliquoted and stored at −80°C until use.

### ELISA

Sera from all rhesus macaques were collected prior to immunization and on weeks 3 and 7 after immunization. Sera was evaluated for SARS-CoV-2 S1-specific IgG using ELISA. ELISA plates were coated with 200 ng of recombinant SARS-CoV-2-S1 protein (Sino Biological) per well overnight at 4°C in carbonate coating buffer (pH 9.5) and then blocked with PBS-T and 2% bovine serum albumin (BSA) for one hour. Rhesus macaque sera was inactivated at 64°C for 40 minutes, then diluted in PBS-T with 1% BSA and incubated overnight. After the plates were washing, anti-monkey IgG-horseradish peroxidase (HRP) (1:50000, Sigma) were added to each well and incubated for one hour. The plates were washed three times, developed with 3,3’5,5’-tetramethylbenzidine, and the reaction was stopped with 1M H_2_SO_4_. Next, absorbance was determined at 450nm using a plate reader (Molecular Devices SPECTRAmax).

### Animals and Immunization

At week 0, male RMs (n=5 animals per group) were bled and primed with 60 μg of tetravalent rS1 proteins of Wuhan, B.1.1.7 (Alpha), B.1.351 (Beta), and P.1 (Gamma) [15μg of each antigen]. Total volume of 300 μl of antigen was mixed with 300 μl of AddaVax adjuvant then administered to RMs (600 μl injection volume). RMs were bled on week 3 and received a homologous booster of 60 μg of tetravalent rS1 proteins. RMs were bled on weeks 7. RMs were also bled and serially euthanized after week 9 post-prime vaccination: on day 0 (RM177), 1 (RM175), 6 (RM176), 8 (RM101), and 15 (RM175). PMBC’s from RMs were collected and analyzed on Days -1, 3, 7, 10, 14, 21, 24, 28, 31, 35, 42, 49, and 64 days post prime immunization. RMs were maintained under specific pathogen-free conditions at the University of Pittsburgh, and all experiments were conducted in accordance with animal use guidelines and protocols approved by the University of Pittsburgh’s Institutional Animal Care and Use (IACUC) Committee.

### SARS-CoV-2 microneutralization assay

Neutralizing antibody (NT-Ab) titers against SARS-CoV-2 were defined according to the following protocol.^80,81^ Briefly, 50 µl of sample from each mouse, starting from 1:10 in a twofold dilution, were added in two wells of a flat bottom tissue culture microtiter plate (COSTAR, Corning Incorporated, NY 14831, USA), mixed with an equal volume of 100 TCID50 of a SARS-CoV-2 Wuhan, Beta, or Delta strain isolated from symptomatic patients, previously titrated, and incubated at 33°C in 5% CO_2_. All dilutions were made in EMEM (Eagle’s Minimum Essential Medium) with addition of 1% penicillin, streptomycin and glutamine and 5 γ/mL of trypsin. After 1 hour incubation at 33°C 5% CO_2_, 3×10^4^ VERO E6 cells [VERO C1008 (Vero 76, clone E6, Vero E6); ATCC® CRL-1586™] were added to each well. After 72 hours of incubation at 33°C 5% CO_2_ wells were stained with Gram’s crystal violet solution (Merck KGaA, 64271 Damstadt, Germany) plus 5% formaldehyde 40% m/v (Carlo ErbaSpA, Arese (MI), Italy) for 30 min. Microtiter plates were then washed in running water. Wells were scored to evaluate the degree of cytopathic effect (CPE) compared to the virus control. Blue staining of wells indicated the presence of neutralizing antibodies. Neutralizing titer was the maximum dilution with the reduction of 90% of CPE. A positive titer was equal or greater than 1:10. The geometric mean titers (GMT) of NT_90_ end point titer were calculated with 4 as a negative shown <10. Sera from mice before vaccine administration were always included in microneutralizaiton (NT) assay as a negative control.

### ACE2 Blocking Assay

Antibodies blocking the binding of SARS-CoV-2 spike variants (Alpha (B.1.1.7), Beta (B.1.351), Gamma (P.1), Delta (B.1.617.2), Zeta (P.2), Kappa (B.1.617.1), New York (B.1.516.1), India (B.1.617 and B.1.617.3)) to ACE2 were detected with a V-PLEX SARS-CoV-2 Panel 18 (ACE2) Kit (Meso Scale Discovery (MSD) according to the manufacturer’s instructions. Antibodies blocking the binding of SARS-CoV-2 spike including Wuhan and spikes from immune evasive variants; BA.1, BA.2, AY.4 (Delta lineage), BA.3, BA.1+R346K mutation, BA.1+L452R mutation, B.1.1.7 (Alpha), B.1.351 (Beta), and B.1.1640.2 to ACE2 were detected with a V-PLEX SARS-CoV-2 Panel 25 (ACE2) Kit (Meso Scale Discovery (MSD) according to the manufacturer’s instructions. Serum samples were diluted (1:10 and 1:100). The assay plate was blocked for 30 min and washed. Serum samples were diluted (1:10 for P18; 1:10 & 1:100 for P25) and 25 μl were transferred to each well. The plate was then incubated at room temperature for 60 min with shaking at 700 rpm, followed by the addition of SULFO-TAG conjugated ACE2, and continued incubation with shaking for 60 min. The plate was washed, 150 μl MSD GOLD Read Buffer B was added to each well, and the plate was read using the QuickPlex SQ 120 Imager. Electrochemiluminescent values (ECL) were generated for each sample. Results were calculated as % inhibition compared to the negative control for the ACE2 inhibition assay, and % inhibition is calculated as follows: % neutralization = 100 × (1 − (sample signal/negative control signal).

### Flow Cytometry

Absolute counts of immune cells in whole blood and immunophenotyping of circulating immune cells were determined by flow cytometry. First, 50 μl of whole blood were added to a TruCount tube (BD Biosciences) containing an antibody mix, allowing to precisely quantify CD45^+^ cell counts in blood, as well as CD4^+^ and CD8^+^ T cells, and CD20^+^ B cells. Whole peripheral blood was stained with fluorescently-labeled antibodies (all purchased from BD Bioscience, San Jose, CA, USA, unless noted otherwise): CD3 (clone SP34-2, V450), CD4 (clone L200, APC), CD8 (clone RPA-T8, PE-CF594), CD28 (clone CD28.2, PE-Cy7), CD38 (clone AT-1, FITC) (Stemcell), CD45 (clone D058-1283, PerCP), CD69 (clone FN50, APC-H7), CD95 (clone DX2, FITC), HLA-DR (clone L243, PE-Cy7), Ki-67 (clone P56, PE). For intracellular staining, cells were fixed and permeabilized with 1X BD Fix/Perm, before being stained for Ki-67. Flow cytometry acquisitions were performed on an LSRFortessa flow cytometer (BD Biosciences), and flow data were analyzed using FlowJo® v10.8.0 (TreeStar, Ashland, OR, USA).

### Spike-Specific Intracellular Staining

Antigen-specific T-cell responses in the PBMC’s of RMs immunized as described above were analyzed after immunization by flow cytometry, adhering to the recently published guidelines.^21,82^ PBMCs collected prior to immunization and on Day 42 post prime immunization were stimulated with PepTivator SARS-CoV-2-S1 (a pool of S1 MHC class I– and MHC class II– restricted peptides) overnight in the presence of protein transport inhibitors (Golgi Stop) for the last 4 hours. Unstimulated cells were used as negative controls. Phorbol myristate acetate (PMA) and ionomycin stimulated cells served as positive controls. Cell were washed with FACS buffer (PBS, 2 % FCS), incubated with Fc Block (BD Biosciences, 553142) for 5 min at 4°C, and stained with surface marker antibody (Ab) stain for 20 min at 4°C. Surface Abs were used as follows: CD3-V450 (SP34-2, V450, BD Biosciences), CD4-APC (L200, APC, BD Biosciences), and CD8ab-PE-CF594 (RPA-T8, PE-CF594, BD Biosciences). For dead cell exclusion, cells were stained with Zombie NIR Fixable Viability dye (BioLegend) for 10 min at 4°C and washed in FACS buffer. Intracellular cytokine staining (ICS) was performed on surface Ab-stained cells by first fixing and permeabilizing cells using the FoxP3 Transcription Factor Staining Buffer kit (eBioscience, 00-5523-00) following manufacturer’s instructions. Intracellular staining with IFNγ-FITC (4S.B3, FITC, BD Biosciences), IL2-PE (MQ1-17H12, PE, BD Biosciences), and TNFa-AF700 (Mab11, AF700, BD Biosciences). Samples were run on an Aurora (Cytek) flow cytometer and flow data were analyzed using FlowJo® v10.8.0 (TreeStar, Ashland, OR, USA).

### Statistical Analysis

Statistical analyses were performed using GraphPad Prism v9 (San Diego, CA). Significant differences are indicated by * p < 0.05. Comparisons with non-significant differences are not indicated.

## Acknowledgements

This work was supported by the National Institutes of Health/National Institute of Diabetes and Digestive and Kidney Diseases/National Institute of Allergy and Infectious Diseases grants R01 DK119936 (CA), R01 DK131476 (CA), R01 AI119346 (CA), R01 DK130481 (IP), R01 DK113919 (IP/CA). AJK was supported in part by the NIH Pitt AIDS Research Training (PART) grant (AI065380) and by the NIAID T32 grant Immunology of Infectious Diseases (IID) (AI060525). AG is funded by NIH grants (UM1-AI106701, R01DK119936-S1 and U01-CA233085) and UPMC Enterprises IPA 25565. The funders had no role in study design, data collection and analysis, decision to publish, or preparation of the manuscript.

## Disclosure

The authors declare that they have competing interests in relation to the research presented in this manuscript. AG, EK, and MSK are co-founders of GAPHAS PHARMACEUTICAL INC., a private startup company that may potentially benefit from the findings of this research. AG, EK, and MSK have equity in GAPHAS PHARMACEUTICAL INC. However, the authors have taken measures to ensure that the research is conducted objectively and that the data and conclusions presented in this manuscript are not influenced by their competing interests. The study was designed, conducted, and analyzed independently of the company. The authors also declare that GAPHAS PHARMACEUTICAL INC. did not provide financial or material support for this research.

---

**Supplementary Figure 1.**
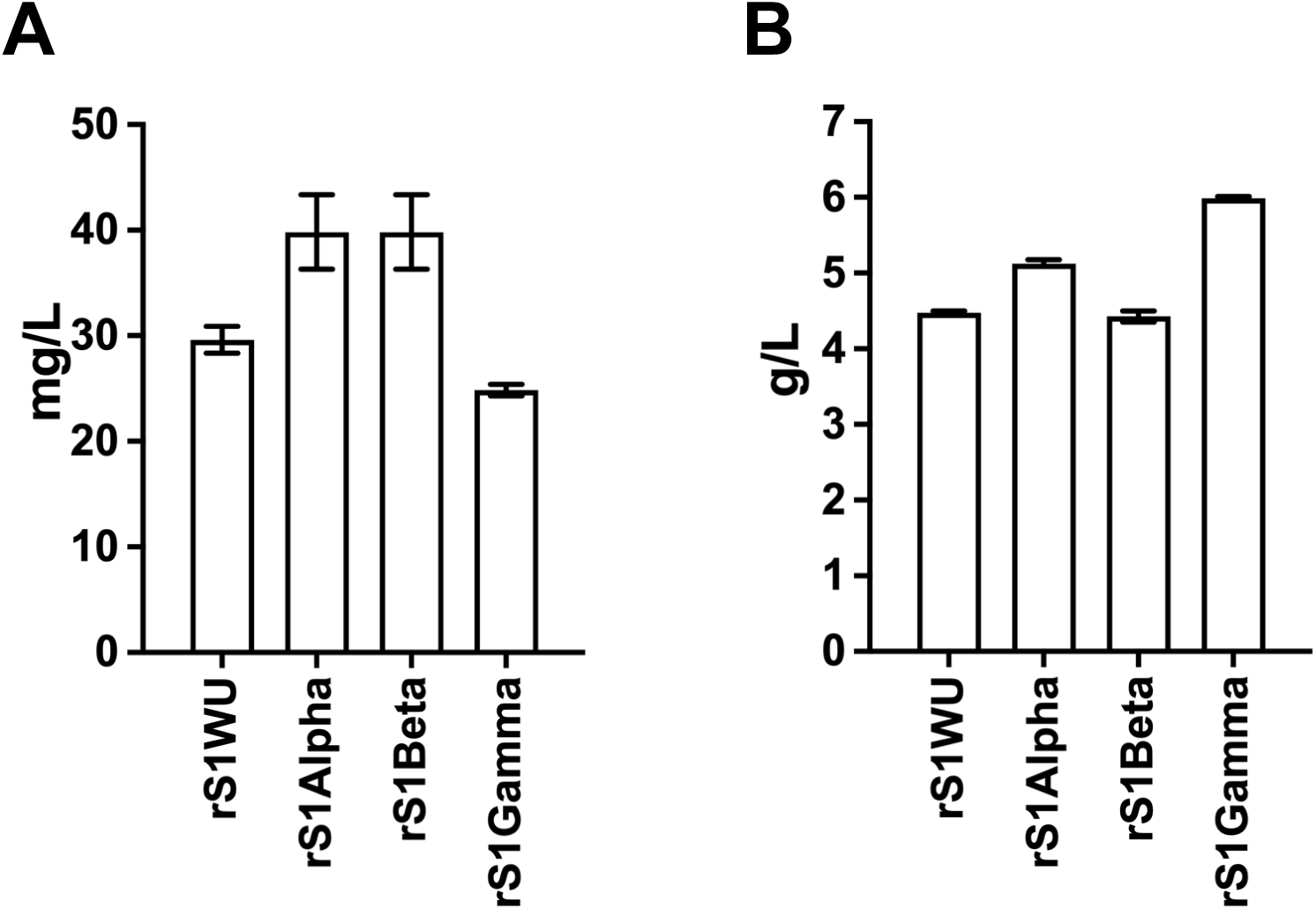
Yield pre and post C-tag purification of each recombinant proteins after transient transfection. To evaluate the expression of rS1WU, rS1Alpha, rS1Beta, and rS1Gamma recombinant proteins, ELISA plates were coated with chimeric MAb 40150-D003 (1:750, Sino Biological) overnight at 4°C. **A.** The supernatants of Expi293^TM^ cells transfected with pAd/S1WU, pAd/S1Alpha, pAd/S1Beta, and pAd/S1Gammawas, respectively, diluted 1:40 or **B.** purified each protein by a CaptureSelect^TM^ C-tagXL Affinity Matrix prepacked column diluted 1:1000 in PBS-T with 1% BSA and along with each purified rS1 proteins for a standard curve were incubated overnight at 4°C. After the plates were washed, chimeric MAb 40150-D001 HRP conjugated secondary antibody (1:10000, Sino Biological) was added to each well. After the development with reagent, the reaction was determined using an ELISA reader (Molecular Devices SPECTRAmax) in same as described in materials and methods.

**Supplementary Figure 2.**
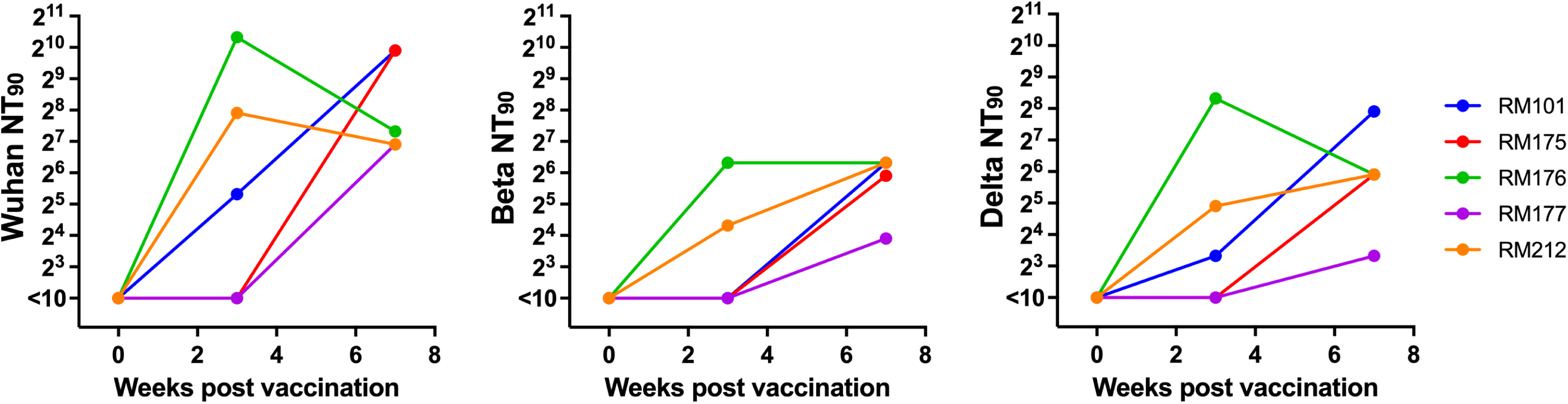
Neutralizing antibodies at week 0, 3, and 7 using a microneutralization assay (NT_90_) were showed in each RM with SARS-CoV-2 Wuhan, Beta, and Delta variants.

