## Supplementary figures and images for "Tetravalent SARS-CoV-2 S1 Subunit Protein Vaccination Elicits Robust Humoral and Cellular Immune Responses in SIV-Infected Rhesus Macaque Controllers"

### Supplementary Figures

#### Slide 1
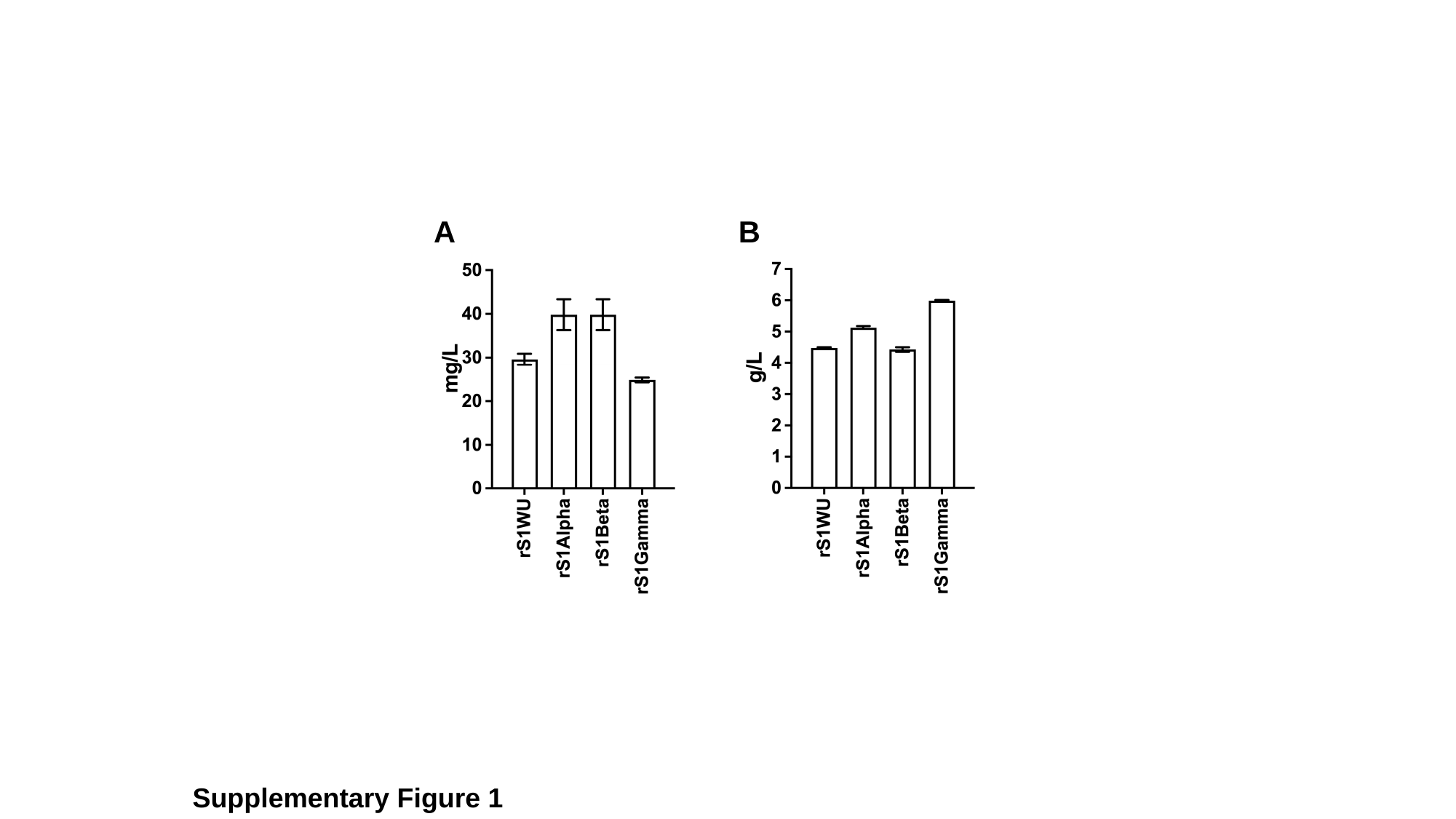

A
B
Supplementary Figure 1

#### Slide 2
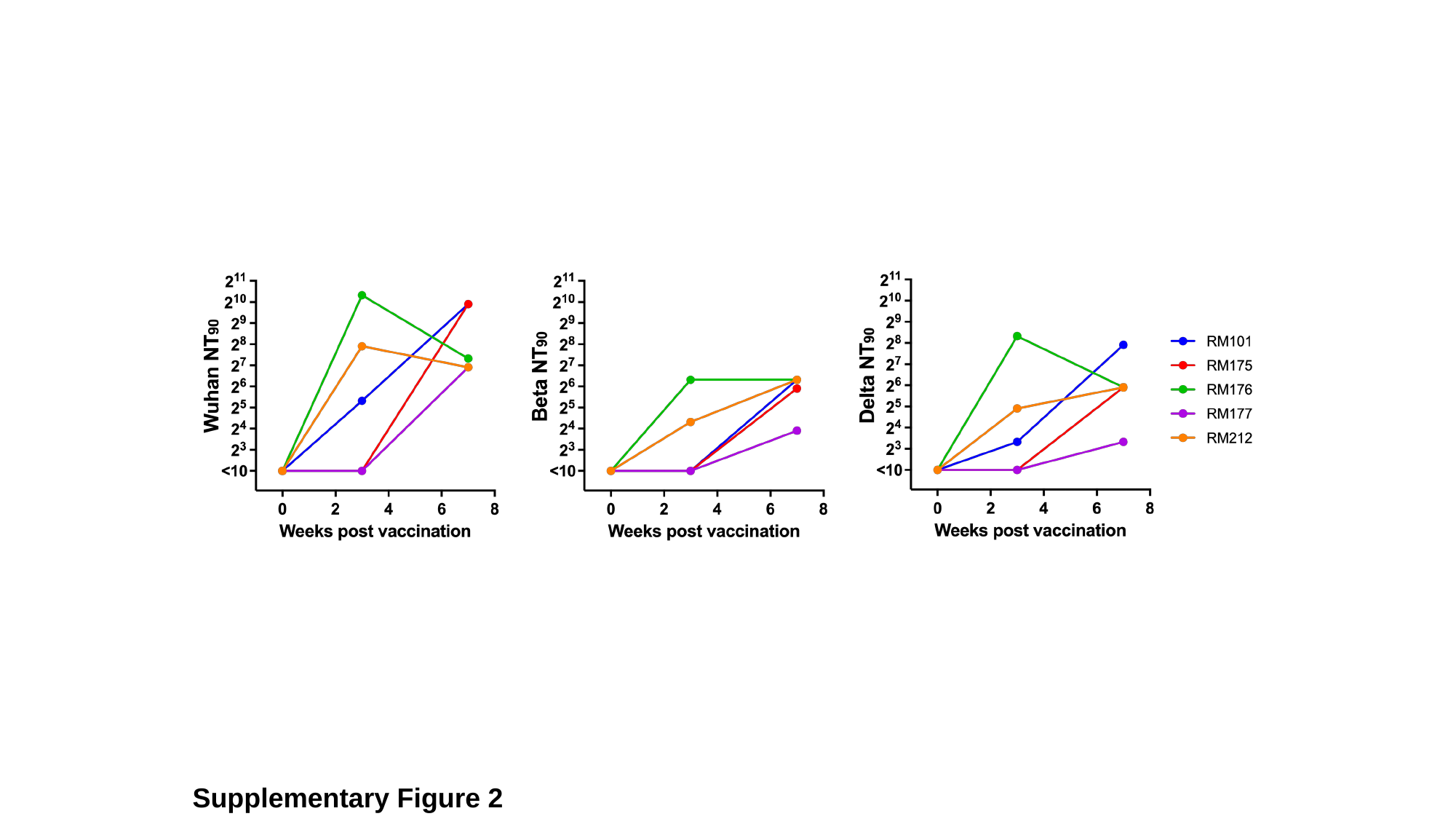

Supplementary Figure 2
